# Heterotrimeric G proteins containing Gβ_1_γ_2_ subunits exhibit subtype-specific mobility differences in live cells

**DOI:** 10.64898/2026.04.13.718213

**Authors:** Ondřej Kuchynka, Andrei Kovalchuk, Mia Nußbaumer, Ekaterina Sviridova, Tomáš Fessl, Alexey Bondar

## Abstract

Heterotrimeric G proteins are key signal transducers in all eukaryotic cells. Anchored to the inner leaflet of the plasma membrane, they transmit and amplify extracellular chemical and physical signals downstream of G protein-coupled receptors. Despite numerous available studies, many biophysical aspects that regulate G protein signaling, including membrane mobility, remain poorly understood. Here, using single-molecule imaging, we show that different subtypes of heterotrimeric G proteins containing Gβ_1_γ_2_ subunits exhibit high diversity in their membrane mobility. We demonstrate that the nature of the Gα subunit plays a major role in defining the mobility of such heterotrimers. Our results indicate that heterotrimers containing Gα_12_ and Gα_13_ subunits exhibit markedly reduced mobility compared to those containing Gα_i/o_ and Gα_s_ subunits. These findings identify subtype-specific lateral membrane mobility of G proteins as a factor that can affect their signaling dynamics in living cells.

## Introduction

G proteins are essential components of cellular signaling. They act as signal transducers in one of the most important and prevalent signaling cascades in the human body, carrying the signal from transmembrane G protein-coupled receptors (GPCRs) to intracellular effectors that regulate the production of secondary messengers. The G protein signaling cascade is involved in nearly every physiological process in humans, and its malfunction is a driving factor behind many diseases, including schizophrenia, heart failure, and cancer [1–3]. Over four decades of research have established the G protein signaling cascade as a target for more than 30% of the drugs available on the market today [4]. While the canonical mechanism of this cascade has been well described, the wider picture, including non-canonical interactions and spatiotemporal signaling dynamics, is only now being uncovered [5].

### G protein localization

Membrane localization of G proteins depends on lipid anchors that ensure their accessibility to receptors and effectors (Fig 1). GPCRs and G proteins alike show an affinity for specific membrane architectures [6]. GPCRs are often surrounded by non-lamellar structures such as hexagonal-phase regions, and some findings suggest that GPCRs might even induce their formation [7]. The affinity of G protein heterotrimers for specific membrane structures has been shown to vary based on lipid anchor types and terminal domain structures of individual G protein subunits [8, 9, 6]. In Gα_i1_β_1_γ_2_ heterotrimers specifically, myristoyl and geranylgeranyl moieties were shown to increase affinity for phosphatidylserine/phosphatidylethanolamine-rich membranes with propensity for non-lamellar structures [8]. At the same time, palmitoylation of Gα_i1_ is thought to induce conformational changes in the subunit, altering the orientation of the G protein’s membrane-facing residues and increasing its affinity for lamellar-prone membranes [10]. Aside from affinity for specific membrane lipid composition, both GPCRs and G proteins also show affinity for particular membrane microdomains, such as lipid rafts. Still, not all G protein subunits seem to share the same microdomain affinity [9, 6]. While activated Gα_s_ and Gα_i1_ are found predominantly in planar lipid rafts, they also populate caveolae in smaller numbers [6]. Activated Gα_q_, on the other hand, shows a strong affinity for caveolae, likely due to an interaction with caveolin [11], and only populates planar lipid rafts in marginal numbers [6]. Similarly, Gβγ subunit dimers also show preferences for specific membrane microdomains based on differences in their prenyl moieties and membrane-facing motifs [12, 13]. A phenomenon linked to the microdomain localization of G proteins is their ability to travel between different cellular compartments, such as endosomes, endoplasmic reticulum, and the Golgi apparatus [14, 15]. The reversible intracellular trafficking of G proteins occurs in heterotrimers and activated Gα and Gβγ alike, and has been linked to non-canonical signal transduction from various organelles, directly connecting microdomain localization of G proteins to signaling regulation [16, 17]. Translocation of G proteins may be connected to various vesicle-mediated endocytic pathways, including clathrin-independent endocytosis through lipid rafts [18, 19, 20], but might also be diffusion-mediated [21]. Overall, G protein localization in membrane domains serves as a key regulator of their functional activity.

**Fig 1.**
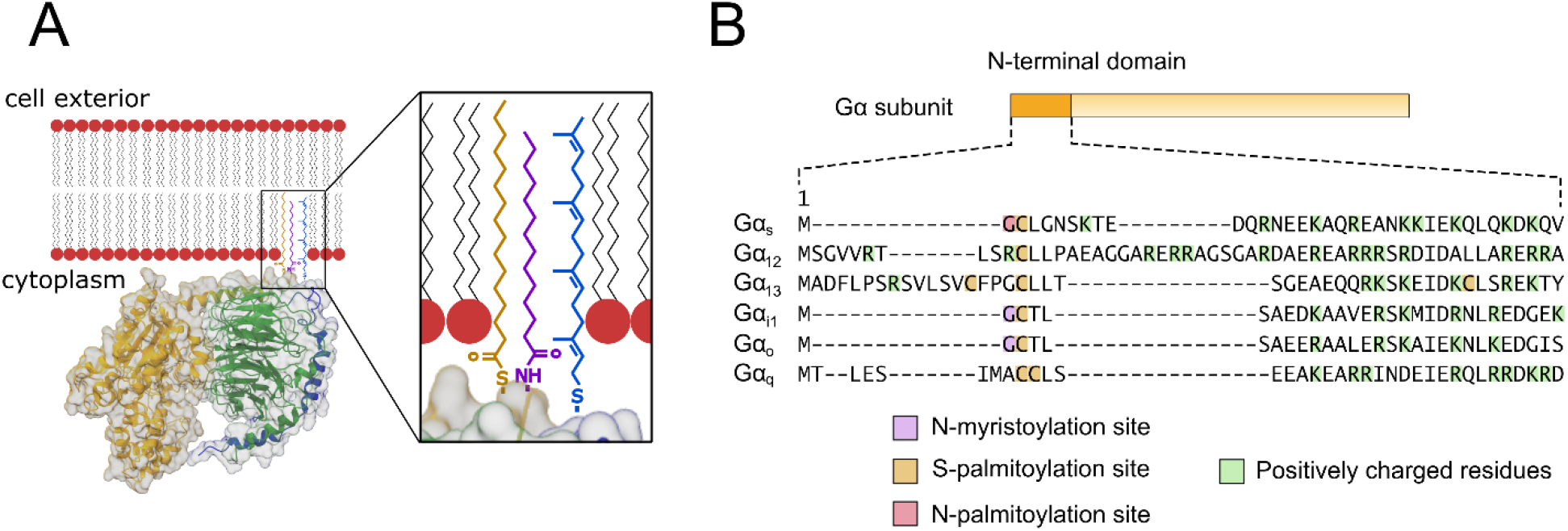
G protein membrane attachment. A) Diagram showing a 3D model of a G protein heterotrimer with subunits Gα (orange), Gβ (green), and Gγ (blue), attached to a cell membrane through lipid moieties — S-geranylgeranyl (blue), N-myristoyl (purple), and S-palmitoyl (orange). B) Diagram showing distinct lipidation sites, as well as the occurrence of positively charged residues in the N-terminal region of different Gα subunits.

### G protein mobility

The membrane environment allows for a wide spectrum of dynamic regulatory mechanisms due to its complexity as a trafficking interface. Mechanistically, the mobility of membrane proteins can be restricted by a system of “fences and pickets”, created by cytoskeletal structures and proteins associated with them [22]. Within this system, protein mobility is further regulated through interactions with the membrane itself and its different microdomains, but also through interactions with other proteins within the microdomains. It has been previously shown that G proteins may exhibit heterogeneous mobility in living cells [23–25]. This heterogeneity was primarily attributed to their interaction with GPCRs in the inactive state (preassembly [26] and inverse coupling [27]) and the active state [25]. The binding of G proteins to the inactive GPCRs leads to a profound decrease in G protein mobility [28, 24]. In contrast, GPCR activation triggers a complex cascade of events, including dissociation of G protein subunits, their interaction with effectors, and relocation to plasma membrane domains, such as clathrin-coated structures and caveolae [25, 23, 29]. A combination of these factors determines the net change in G protein mobility upon activation. In addition to signaling interactions, the cytoskeleton, and actin in particular, has been implicated in forming a grid that confines the movement of G proteins to individual compartments and enables spatial restriction of signaling propagation [24, 25]. The role of membrane nanodomains is also related to the confinement of G protein localization, particularly after dissociation of Gα and Gβγ subunits. Beyond the interactions with membrane proteins and microdomains, significant differences in G protein mobility based on both the Gα and Gγ subunit subtype composition have been reported [30, 12]. However, despite existing evidence of composition-based dynamic variability in G proteins, the diversity and scope of this effect remain insufficiently understood.

Here, we set out to determine whether G protein heterotrimers containing Gβ_1_γ_2_ subunits and different subtypes of Gα subunit exhibit distinct mobility patterns in living cells. Using single-molecule imaging, we demonstrate that Gα subunit identity plays an important role in the mobility of these heterotrimeric G proteins.

## Results

### Determination of G protein mobility by single-particle tracking

We utilized Total Internal Reflection Fluorescence (TIRF) microscopy and single-particle tracking to determine the mobility patterns of different classes of G proteins in the plasma membrane of living cells (Fig 2A, 2B). Since direct labeling of Gα subunits has been shown to affect their mode of interaction with Gβγ [31, 32], we relied on labeling the Gγ_2_ subunit, which we used as a universal readout sensor for mobility comparison of different classes of G proteins. In our experiments, CHO-K1 cells were transiently transfected with non-tagged Gα and Gβ_1_ subunits overexpressed under the CMV promoter, in combination with the Gγ_2_-HaloTag subunit expressed at a low level (0.14-0.20 molecules/μm^2^; S5 Fig, S1 Table) using the *minP* promoter (Promega) and efficiently labeled with the Halo-JF646 fluorescent ligand. This arrangement allowed for the majority of Gγ_2_-HaloTag molecules to preferentially localize in heterotrimers containing the studied Gα. We captured single-molecule data as a kinetic series of 1000 frames with a frame rate of 40 FPS at 37°C (representative frames are shown in Fig 2B and S5A-5G Fig). Spot detection and tracking were performed in TrackMate [33] (Fig 2C, 2D), and diffusion states were determined using Divide-and-Conquer Moment Scaling Spectrum (DC-MSS) diffusion analysis [34]. Data summary is shown in supporting Table 2.

**Fig 2.**
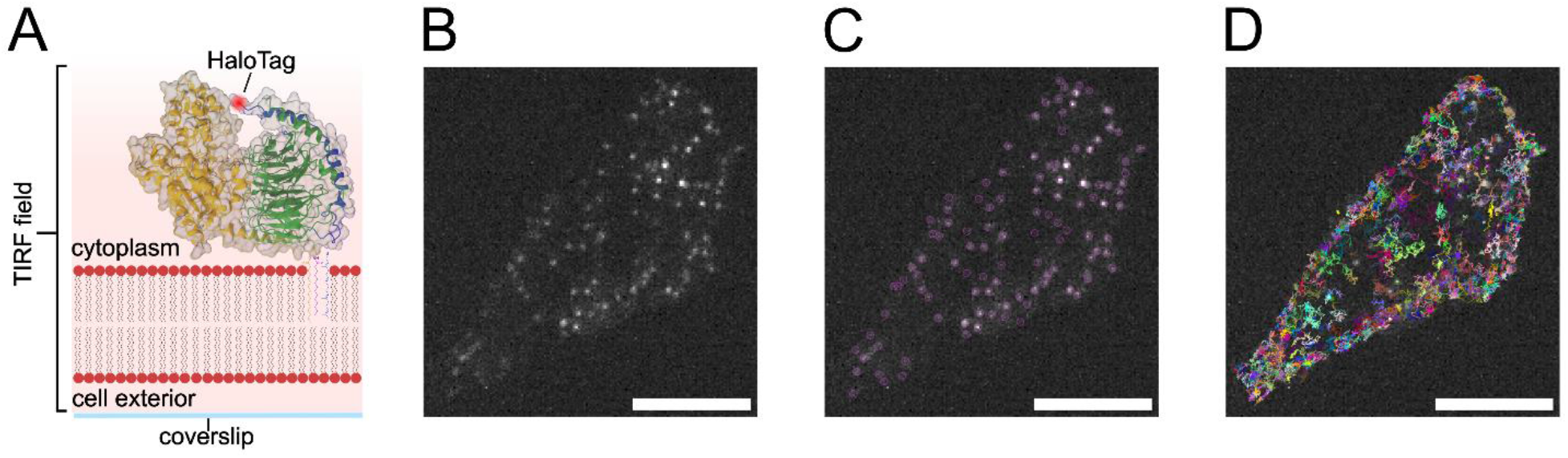
Experimental setup and workflow. A) 3D model of a G protein heterotrimer with subunits Gα (orange), Gβ (green), and Gγ (blue), attached to a cell membrane during TIRF imaging. Positioning of the HaloTag at the N-terminus of Gγ_2_ is highlighted in red. B-D) Images demonstrating the three main steps of image analysis, including raw data from TIRF imaging (B), molecule detection (C), and track generation (D). The scale bar in B-D is 10 µm.

### Different families of G proteins containing Gβ_1_γ_2_ subunits exhibit distinct mobility patterns in live cells

We first assessed the mobility of six representative G proteins belonging to all four families, specifically G_s_, G_i1_, G_o_, G_q_, G_12_, and G_13_, in the plasma membrane of live cells (Fig 3A, 3B) using Gγ_2_-HaloTag as a reporter in all the measurements. Strikingly, we found that G proteins containing Gα_12_ and Gα_13_ subunits showed significantly lower diffusion coefficients (0.14 and 0.20 μm^2^/sec, respectively) than those containing Gα_s_, Gα_i1_, and Gα_o_ subunits (0.30, 0.34, and 0.29 μm^2^/sec, respectively). The G_q_ protein mobility was somewhat in between these groups (0.23 μm^2^/sec). Surprisingly, the observed pattern did not correlate with the number of lipid anchors on the Gα subunit: Gα_12_ only contains a single lipid anchor on its N-terminus, Gα_13_ contains three, and other tested Gα subunits contain two N-terminal lipid anchors (Fig 1B). The differences in mobility could indicate either the overall slower movement of G_12_ and G_13_ compared to other tested G proteins or a difference in the distribution of the diffusion states between these proteins. Slower overall movement is usually characteristic of bulkier proteins or integral membrane proteins, while redistribution of diffusion states can point to interaction with additional molecular partners (e.g., GPCRs).

**Fig 3.**
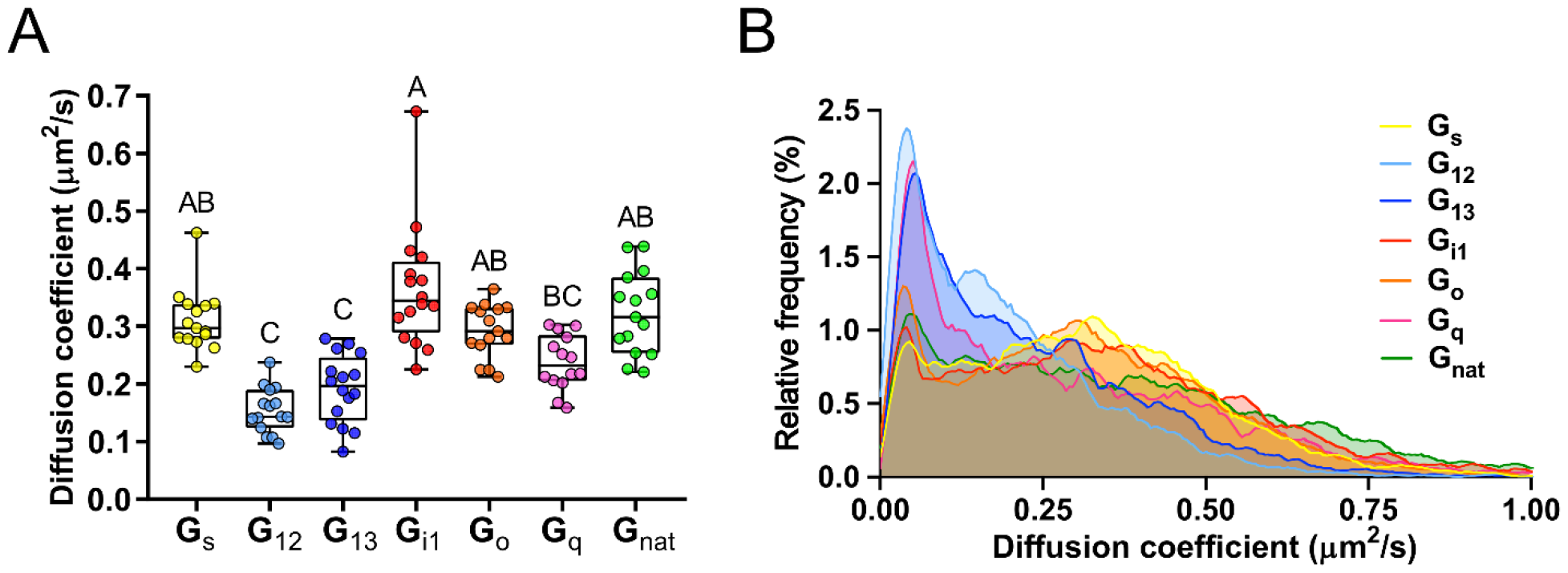
Different G protein families show distinct mobility patterns. A) Box plots of median diffusion coefficients derived from individual cells expressing heterotrimers G_s_ (yellow), G_12_ (light blue), G_13_ (dark blue), G_i1_ (red), G_o_ (orange), G_q_ (magenta), and G_nat_ (green), containing Gβ_1_ subunits with Gγ_2_-HaloTag. The boxes indicate the median and the interquartile range. Letters above individual datasets indicate similarity grouping based on statistical analysis (*α* = 0.05, *p* < 0.0001, *H* = 63.81, *ε^2^* = 0.58). B) Graphical comparison of the distribution of diffusion coefficients of molecule tracks of labeled G_s_ (yellow), G_12_ (light blue), G_13_ (dark blue), G_i1_ (red), G_o_ (orange), G_q_ (magenta), and G_nat_ (green). No statistical inference was performed on these distributions.

### G proteins demonstrate differences in the distribution of mobility states

To further address the differences in mobility of the assessed G proteins, we examined the occurrence and distribution of distinct diffusion states among these G proteins. G proteins containing subunits Gα_12_, Gα_13_, and Gα_q_ exhibited significantly higher mean per-cell fractions of molecules with restricted movement – immobile and confined (59, 59, and 57% respectively), compared to those containing Gα_s_, Gα_i1_, and Gα_o_ subunits (46, 39, and 42% respectively) (Fig 4, S1 and S2 Figs). This bias in diffusion state occurrence partially explains the mobility variance described earlier, showing that G_12_, G_13_, and G_q_ exhibit lower mobility than G_s_, G_i1_, and G_o_. The other aspect of G protein mobility variance we observed is that G proteins containing subunits Gα_12_ and Gα_13_ consistently displayed lower mobility than G proteins containing subunits Gα_s_, Gα_i1_, Gα_o_, and Gα_q_, even within specific diffusion states (S3 Fig). These findings indicate that G_12_ and G_13_ differ from the other G proteins on multiple levels.

**Fig 4.**
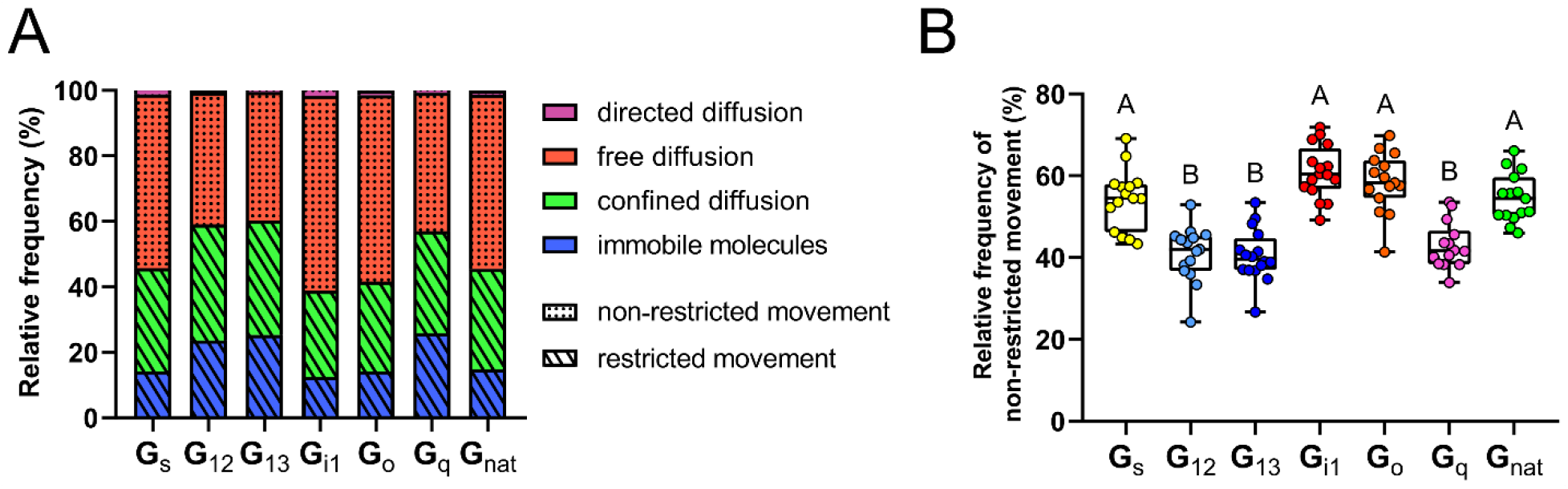
Occurrence of individual diffusion states in labeled G proteins. A) Fractions of immobile molecules (blue) and molecule tracks exhibiting confined (green), free (red), and directed diffusion (purple). Additionally, fractions of molecule tracks exhibiting restricted (immobile molecules and confined diffusion, diagonal line pattern) and non-restricted movement (free and directed diffusion, dotted pattern) are shown. B) Box plots of relative frequency of non-restricted movement in labeled G_s_ (yellow), G_12_ (light blue), G_13_ (dark blue), G_i1_ (red), G_o_ (orange), G_q_ (magenta), and G_nat_ (green). The boxes indicate the median and the interquartile range. Letters above individual datasets indicate similarity grouping based on statistical analysis (*α* = 0.05, *p* < 0.0001, *H* = 67.59, *ε^2^* = 0.62).

We also determined the mobility of fluorescently labeled Gβ_1_γ_2_ subunits expressed in cells without exogenous Gα subunits. The cells were transfected with constructs encoding overexpressed Gβ_1_ subunits and low-expressed Gγ_2_-HaloTag subunits, which were left to pair with endogenous Gα subunits, producing G proteins here referred to as G_nat_. We found that G_nat_ proteins displayed mobility roughly similar to that of G_s_ or G_i1_, both in terms of the diffusion coefficient (0.32 μm^2^/sec) and the occurrence of diffusion states (45% of molecules exhibited restricted movement) (Figs 3 and 4). G_nat_ also presented a uniquely wide distribution of diffusion coefficient values, with very low bias for specific mobility range, especially evident in freely diffusing molecules (S1 and S2 Figs). The wide distribution and lack of bias indicate that G_nat_ possibly incorporates a number of different Gα subunits with different mobility patterns. At the same time, the data show that G proteins containing subunits Gα_s_ and Gα_i1_ exhibit mobility closest to that of the “average” G protein containing an endogenous Gα subunit, making Gα_12_ and Gα_13_ clear outliers with respect to mobility of Gβ_1_γ_2_-containing heterotrimers.

## Discussion

G proteins are molecular messengers in one of the most prevalent signaling cascades in eukaryotes. Their dynamic behavior is a key regulator of signal transfer dynamics [25]. Native biological systems present a complex environment in which mobility can be modulated through protein interactions, localization in membrane nanodomains, or preference towards a particular membrane structure. Surprisingly, we observed profound differences in the mobility of G protein heterotrimers differing only in their Gα subunits, with G_12_ and G_13_ showing significantly lower mobility than G_s_, G_i1_, and G_o_ proteins. The mobility of membrane proteins is closely tied to interactions with their surroundings. G protein signaling naturally involves binding to GPCRs, effectors, regulators, and membrane domains, and the mechanisms of these interactions can vary greatly. While many interactions are transient, some can involve the formation of lasting low-mobility complexes such as the oligomeric precoupling complex of muscarinic receptor M4 and protein G_o_ [35], muscarinic receptor M3 and G_q_ [26], or the inverse coupling of G_s_ to the 5-HT_7_ receptor [27]. Although, to the best of our knowledge, there is no direct evidence of precoupling or inverse coupling for G_12_ or G_13_, undiscovered interactions with a receptor, effector, membrane domain, or the actin cytoskeleton present a possible explanation for some of the observed variance in G protein mobility.

Membrane affinity of Gα subunits plays a vital role in membrane attachment and is connected to the N-terminal architecture and lipidation of individual subunits. However, in our study, the Gα subunits that formed heterotrimers exhibiting lower mobility: Gα_12_, Gα_13_, and Gα_q_ (Fig 3), not only lack a common lipidation motif, but also boast high variability in terms of N-terminal lipidation sites. Gα_13_ carries three distinct S-palmitoylation sites, while Gα_q_ only carries two, and Gα_12_ carries only a single S-palmitoylation site, which is the lowest in any of the tested subunits (Fig 1B). We conclude that the number of Gα subunit lipidation sites is not a major driver of the observed dynamic variability of G proteins, at least in heterotrimers containing Gβ_1_γ_2_ subunits.

Even though Gα_12_, Gα_13_, and Gα_q_ do not share a common lipidation motif, there is one aspect of the N-terminal lipidation that is consistent in all the low-mobility subunits, which is the lack of N-palmitoylation/N-myristoylation. All Gα subunits carry at least one lipid tag in their N-terminal region, but only some undergo a truly N-terminal lipidation. Lipidation in subunits Gα_12_, Gα_13_, and Gα_q_ only occurs further up the N-terminal sequence on positions 9-36 (Fig 1B). While there is no known mechanism that would directly tie lipid anchor linkage to the observed differences in mobility, some studies suggest a differential effect of thioester-linked and amide-linked lipid anchors on G protein dynamics [36]. Partial lipidation may also be a contributing factor of the observed heterogeneity [37]. Ascertaining whether these differences in lipid anchor linkage and dynamics are involved in modulating G protein mobility requires further investigation.

Aside from distinct lipidation patterns, the N-terminal region of the Gα subunit presents two other factors that may contribute to the mobility of G proteins: the size of the N-terminal tail and the size of the positive patch motif. Both of these are particularly pronounced in palmitoylated subunits and could impact the interaction of heterotrimers with the membrane or other proteins. The N-terminal region is by far the largest in subunits Gα_12_ and Gα_13_ and could be involved in effects such as steric hindrance, but the structural properties of this expanded structure have yet to be fully deciphered. The positive patch motif can play a role in the membrane trafficking of newly synthesized Gα subunits along with the N-terminal myristoyl moiety and the Gβγ subcomplex [38]. In a study focused on describing the positive patch architecture across various Gα subunits, Kosloff et al. found that Gα_12_ showed the largest positive patch among the non-myristoylated Gα subunits [38]. While this finding aligns with our results in G_12_ (Fig 3), the article also reports the presence of a sizable positive patch on the N-terminus of Gα_s_, which in our study exhibited mobility comparable to the myristoylated subunits with much smaller positive patches (Fig 3). Another study by Mystek et al. suggests that not only the size of the positive patch but also the nature and exact position of charged amino acids together define the affinity of Gα subunit N-termini for the plasma membrane [39]. The exact role of the Gα N-terminal architecture in G protein mobility, therefore, remains to be elucidated.

In conclusion, our results demonstrate strong differences in mobility of different types of heterotrimeric G proteins containing Gβ_1_γ_2_ subunits in live cells and point to a potentially underappreciated mechanism of G protein signaling cascade regulation that utilizes the localization and accessibility of G protein molecules rather than direct regulation of their conformation.

## Limitations

In this study, we present evidence of dynamic variability in G protein heterotrimers containing Gγ_2_-HaloTag, Gβ_1_, and various Gα subunits, offering a closer look at the previously ill-described dynamic behavior of these molecular messengers. At the same time, several limitations remain within the study.

Firstly, our fluorescent reporter design did not allow for direct evaluation of the expression levels of individual Gα subunits in the measured cells. However, direct labeling of the Gα_s_ subunit with HaloTag yielded a diffusion coefficient similar to that observed in the G_s_ experiments with Gγ_2_-HaloTag (S6A Fig). Furthermore, activation of G_s_ labeled with Gγ_2_-HaloTag using vasopressin receptor type 2 (V2R) led to changes in the diffusion coefficient consistent with those previously observed for G_s_ activation by β_2_-adrenoceptor [23], indicating the formation of functional G_s_ heterotrimers containing Gγ_2_-HaloTag (S6B Fig). Additionally, experiments with heterotrimers containing mutant Gα_s_ lacking two C-terminal amino acids (G_s_-ΔCt2) produced higher diffusion coefficients in Gγ_2_-HaloTag, indicating fewer interactions with slow membrane-localized binding partners (GPCRs) (S6A Fig). Nevertheless, expression level differences remain a potential source of additional variability within our dataset.

Secondly, the universal Gγ_2_-HaloTag readout sensor used in our study does not report the mobility of individual Gα subunits directly, but rather through their pairing partners. This approach does not allow us to determine the exact fraction of labeled Gβ_1_γ_2_ associated with the exogenously expressed Gα subtype, and it could introduce external variability in the event of heterotrimer misassembly. At the same time, G_s_ heterotrimers containing Gγ_2_-HaloTag seemingly respond to V2R activation (S6B Fig) and exhibit mobility similar to that of G_s_ heterotrimers labeled at the Gα subunit (S6A Fig). These data suggest that our strategy yields functional heterotrimers that predominantly contain the exogenous Gα subunits. Additionally, the mobility observed in G_nat,_ which closely resembles the scenario of complete exogenous Gα misassembly, sharply diverges from that observed in G_12_ and G_13_ (Fig 3B), indicating that the observed variance cannot be explained solely by differences in heterotrimer assembly.

Thirdly, the density of expressed molecules may affect diffusion coefficients. Our linear regression analysis shows no correlation between the membrane density of Gγ_2_-HaloTag and observed diffusion coefficients (S5I Fig). However, potential variations in the expression of Gα and Gβ impose limits on resulting data interpretation.

Fourthly, the physiological relevance of the quantitative values for diffusion coefficients has limited value. Indeed, our experiments in HEK293 cells demonstrate that G_s_ diffusion in these cells is different from that measured in CHO-K1 cells (S6 Fig). We believe that any absolute values of this kind will have limited significance; however, the differences between the values are valuable and highlight important trends in the behavior of G protein heterotrimers.

Furthermore, the focus of the current study is limited to examining the effect of the Gα subunit on the mobility of heterotrimers containing the Gγ_2_ subunit. To generalize G protein mobility dynamics beyond this point, additional subunit combinations need to be evaluated, particularly those with different subtypes of both Gα and Gγ subunits.

## Materials and Methods

### DNA constructs

The constructs Gα_s_, Gα_i1_, Gα_12_, Gα_o_, and Gβ_1_ in pcDNA3.1 were purchased from the cDNA Resource Center (cdna.org). Gα_13_-IRES-mCherry and Gα_q_-IRES-mCherry constructs were generously provided by Dr. Josef Lazar (Charles University, Prague). SNAPf-V2R and Gα_s_-ΔCt2 constructs were provided by Nevin A. Lambert (Augusta University). minP-HaloTag-Gγ_2_ and Gα_s_-HaloTag constructs were published previously [23].

### Cell culture and transfection

CHO-K1 cells (85051005 ECACC) and HEK293 cells (CRL-1573 ATCC) were cultured in F-12K and DMEM medium, respectively, supplemented with 10% Fetal Bovine Serum and an antibiotic/antimycotic mixture at 37 °C and 5% CO_2_. Cells were regularly tested for the absence of mycoplasma contamination. Transient transfection was performed using Lipofectamine 3000 (ThermoFisher) according to the manufacturer’s instructions. All experiments were conducted two days post-transfection.

### Sample preparation for TIRF microscopy

Before imaging, the transfected cells were labeled with the Halo-JF646 fluorescent dye (Promega). The cells were covered with 1 ml of supplemented F-12K/DMEM medium mixed with 2 µl of 50 nM Halo-JF646 and incubated for 20-30 minutes to allow for proper labeling.

After labeling, the cells were washed three times with 750 µl of DPBS −/−. The cells were then dislodged by adding 500 µl of Accutase (Merck) for 8-12 minutes at room temperature. 500 µl of DPBS +/+ was added, and the resulting suspension was mixed, transferred to a 1.5 ml Eppendorf tube, and centrifuged at 2900 RPM for 3 minutes. After the spin, the supernatant was removed, and 1 ml of DPBS +/+ was added and mixed to break apart the cell pellet. This washing step was repeated twice, with the tube spun at 2900 RPM for 2 minutes, first in DPBS and then in SFM4CHO medium. After the third spin, the SFM4CHO medium supernatant was removed and replaced with 1 ml of fresh SFM4CHO, creating the final cell suspension.

Ultra-clean coverslips were first removed from their ethanol container and placed in culturing wells, with the coverslips leaning against the well wall to allow the ethanol to evaporate. After all the ethanol had evaporated, the coverslips were properly rested onto the bottom of the well and covered with a solution of 5 µl of fibronectin and 500 µl of DPBS +/+. The coverslips were left to rest at room temperature for at least 30 minutes. The fibronectin was washed off by adding 1 ml of DPBS +/+ and removing 1 ml of the solution, which was repeated 3 times. Finally, all the remaining DPBS solution was removed, and the labeled cell suspension was transferred onto the coverslip. The amount of cell suspension used was adjusted based on prior cell confluence to 0.2-1 ml, accompanied by 1-1.8 ml of SFM4CHO medium. After that, the wells were incubated for at least an hour to allow for proper attachment to the coverslip. Before imaging, the coverslip with cells was washed twice with 1 ml of DPBS +/+ and transferred into an Attofluor cell chamber (Invitrogen).

### Single-molecule TIRF microscopy

TIRF microscopy was performed on the inverted RAMM microscope (ASI imaging) equipped with a 60× 1.49 NA objective lens (Olympus) and an iChrome CLE laser light source (Toptica) (S4 Fig). Emitted fluorescence was filtered using an FF640-FD:01 dichroic beamsplitter (Semrock) and a Brightline 661/20 filter (Semrock). The data were captured as a kinetic series of 1000 frames with a frame rate of 40 FPS at 100% laser power. The image series were obtained in a single channel with a 640 nm laser (20mW). Cell selection was based primarily on the molecular membrane density of labeled G proteins (∼ 0.1-0.3 molecules/µm^2^) and was conducted at 10 FPS and 30% laser power to reduce photobleaching. To keep experiments consistent, the mCherry signal from Gα_13_ and Gα_q_ constructs was not used as a selection criterion. The data were acquired using the SONA sCMOS camera (Andor). All experiments were done at 37°C using an environmental chamber (OkoLab). The imaging data from this study are available at the EMBL-EBI BioImage Archive (doi:10.6019/S-BIAD3061).

### Image analysis

Image analysis was carried out in two parts. TrackMate ImageJ plugin was used for single-molecule detection and molecule track generation [33]. Then, a MATLAB-based DC-MSS analysis by Vega et al. [34] was used for diffusion coefficient determination and diffusion state characterization.

In Trackmate, molecules in individual time series were detected using the LoG detector with subpixel localization precision, and the simple LAP tracker was used to track their movement across the series and create track records. For spot detection, the suitable particle size of our molecules was determined to be 0.6 µm and was consistently used for all samples throughout the analysis. The spot detection was further refined through the built-in TrackMate quality thresholding on a case-by-case basis to maximize the detection accuracy. During the subsequent track generation, the linking distance was adjusted in the range of 0.6-1 µm, reflecting the sample mobility variability. The value of gap closing distance was also adjusted according to the sample in the range of 0.8-1.2 µm. The number of gap frames was kept at 2 to allow tracks to be reconnected only in the case of a single missing frame, preventing the conflation of unrelated molecule tracks. The tracks were filtered to remove tracking artifacts using the Mean Directional Change Rate and Maximum Straight Line Speed thresholding. The three major steps of the TrackMate analysis are showcased in Fig 2B-2D.

To assess the robustness of our data analysis approach, we analyzed a representative dataset using an alternative approach with fixed input values of quality thresholding = 4, linking distance = 0.7, and gap closing distance = 0.8, optimized to avoid inaccurate track splitting and merging across conditions. The results show that significant differences in the mobility of G_s_ and G_12_ persist, regardless of the analysis approach (S7 Fig).

An in-house-built MATLAB script incorporating the DC-MSS analysis was designed for the purposes of molecule track analysis. The first step in the analysis was reformatting the XML output data from TrackMate into a MATLAB format suitable as DC-MSS input. This data was then subjected to the DC-MSS analysis as described by Vega et al. [34]. The DC-MSS analysis works to detect motion type switches upon which it generates track segmentation. The tracks are then classified based on MSS analysis of molecule displacements, in which segments are assigned one of the following four diffusion states: immobile molecules, confined diffusion, free diffusion, and directed diffusion. Lastly, the classified segments are subjected to adjacent merging and reclassification. Merges are accepted only if the reclassified merged segment is of the same or lower mobility class. This approach enables the detection of a large number of switch events while also preventing overall oversegmentation through two separate merging steps. After the DC-MSS analysis was complete, we extracted segment information, including diffusion coefficient value, segment length, and diffusion state, and calculated median values and diffusion state populations for each cell. Only segments longer than 500ms were included in the results. The full code is available in a public repository (https://doi.org/10.5281/zenodo.21722781).

### Statistical analysis

Kruskal-Wallis test followed by Dunn’s multiple comparisons test with adjusted p-values controlling the family-wise error rate was used to evaluate the significance of our results (Figs 3A and 4B; S3A-3D Fig). The built-in GraphPad Prism multiple comparisons correction was used during the analysis. The reason for using the Kruskal-Wallis test over one-way ANOVA was the non-normal distribution and occasional inequality of variances in our datasets. The observed effect size was calculated using the epsilon squared (ε^2^) statistic using the following formula.

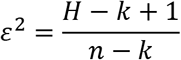

Where *H* is the Kruskal-Wallis test statistic, *k* is the number of groups, and *n* is the total sample size.

For pairwise comparisons (S6C and S7 Figs), a two-sided Mann-Whitney U test computing exact P values was used instead. The observed effect size was calculated using the rank-biserial correlation (r_rb_) statistic, as follows.

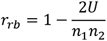

Where *U* is the Mann-Whitney test statistic, and *n_1_* and *n_2_* are the sample sizes of each group.

Ordinary least squares regression was used to evaluate the relationship between the membrane density of Gγ_2_-HaloTag (*x*) and the median diffusion coefficients (*y*) of each condition. The regression line was fitted without constraining the intercept, and 95% confidence intervals were calculated. Statistical tests were two-sided.

Due to the limited number of observations per condition, regression lines fitted separately for each experimental group were used for visualization only and were not subjected to statistical inference (S5H Fig). Formal assessment of the association between membrane density and median diffusion coefficients, while accounting for group differences, was performed by linear regression of residuals obtained after mean-centering both variables within each condition (S5I Fig).

Statistical analysis was carried out in GraphPad Prism 8.0.1. A priori power analysis was conducted using G*Power 3.1.9.7, using the one-way fixed-effects ANOVA model as an approximation for the Kruskal-Wallis test. A large effect size (f = 0.40) was assumed based on preliminary data.

The statistical unit used for analysis was cell-level medians. Individual molecule tracks were treated as descriptive observations, used only for graphical representation. No statistical inference was performed on track-level values.

## Acknowledgements

We thank Prof Nevin A Lambert (Augusta University) for inspiration, helpful discussions, and provision of constructs. We thank Paul Miclea and Josef Lazar (Charles University) for the provision of constructs.

## References

1. Boczek T, Mackiewicz J, Sobolczyk M, Wawrzyniak J, Lisek M, Ferenc B, et al. The Role of G Protein-Coupled Receptors (GPCRs) and Calcium Signaling in Schizophrenia. Focus on GPCRs Activated by Neurotransmitters and Chemokines. Cells. 2021;10: 1228. doi:10.3390/cells10051228

2. Wang J, Gareri C, Rockman HA. G-Protein–Coupled Receptors in Heart Disease. Circulation Research. 2018;123: 716–735. doi:10.1161/CIRCRESAHA.118.311403

3. Yu S, Sun L, Jiao Y, Lee LTO. The Role of G Protein-coupled Receptor Kinases in Cancer. Int J Biol Sci. 2018;14: 189–203. doi:10.7150/ijbs.22896

4. Caroli J, Andreassen SN, Lorente JS, Xiao B, Pándy-Szekeres G, Gloriam DE. An online GPCR drug discovery resource. npj Drug Discov. 2025;2: 17. doi:10.1038/s44386-025-00010-9

5. Bondar A, Lazar J. Optical sensors of heterotrimeric G protein signaling. The FEBS Journal. 2021;288: 2570–2584. doi:10.1111/febs.15655

6. Oh P, Schnitzer JE. Segregation of Heterotrimeric G Proteins in Cell Surface Microdomains. MBoC. 2001;12: 685–698. doi:10.1091/mbc.12.3.685

7. van der Wel PCA, Pott T, Morein S, Greathouse DV, Koeppe RE, Killian JA. Tryptophan-Anchored Transmembrane Peptides Promote Formation of Nonlamellar Phases in Phosphatidylethanolamine Model Membranes in a Mismatch-Dependent Manner. Biochemistry. 2000;39: 3124–3133. doi:10.1021/bi9922594

8. Álvarez R, Escribá PV. Structural Basis of the Interaction of the G Proteins, Gαi1, Gβ_1_γ_2_ and Gαi1β1γ2, with Membrane Microdomains and Their Relationship to Cell Localization and Activity. Biomedicines. 2023;11: 557. doi:10.3390/biomedicines11020557

9. Waheed AA, Jones TLZ. Hsp90 Interactions and Acylation Target the G Protein Gα12 but Not Gα13 to Lipid Rafts. Journal of Biological Chemistry. 2002;277: 32409–32412. doi:10.1074/jbc.C200383200

10. Álvarez R, López DJ, Casas J, Lladó V, Higuera M, Nagy T, et al. G protein–membrane interactions I: Gαi1 myristoyl and palmitoyl modifications in protein–lipid interactions and its implications in membrane microdomain localization. Biochimica et Biophysica Acta (BBA) - Molecular and Cell Biology of Lipids. 2015;1851: 1511–1520. doi:10.1016/j.bbalip.2015.08.001

11. Calizo RC, Scarlata S. A Role for G-Proteins in Directing G-Protein-Coupled Receptor– Caveolae Localization. Biochemistry. 2012;51: 9513–9523. doi:10.1021/bi301107p

12. Abankwa D, Vogel H. A FRET map of membrane anchors suggests distinct microdomains of heterotrimeric G proteins. J Cell Sci. 2007;120: 2953–2962. doi:10.1242/jcs.001404

13. Mystek P, Rysiewicz B, Gregrowicz J, Dziedzicka-Wasylewska M, Polit A. Gγ and Gα Identity Dictate a G-Protein Heterotrimer Plasma Membrane Targeting. Cells. 2019;8: 1246. doi:10.3390/cells8101246

14. Khater M, Wei Z, Xu X, Huang W, Lokeshwar BL, Lambert NA, et al. G protein βγ translocation to the Golgi apparatus activates MAPK via p110γ-p101 heterodimers. J Biol Chem. 2021;296: 2191–2198. doi:10.1016/j.jbc.2021.100325

15. Yu J-Z, Rasenick MM. Real-Time Visualization of a Fluorescent Gαs: Dissociation of the Activated G Protein from Plasma Membrane. Molecular Pharmacology. 2002;61: 352–359. doi:10.1016/S0026-895X(24)12868-4

16. Kankanamge D, Tennakoon M, Karunarathne A, Gautam N. G protein gamma subunit, a hidden master regulator of GPCR signaling. Journal of Biological Chemistry. 2022;298. doi:10.1016/j.jbc.2022.102618

17. Saini DK, Chisari M, Gautam N. Shuttling and translocation of heterotrimeric G proteins and Ras. Trends in Pharmacological Sciences. 2009;30: 278–286. doi:10.1016/j.tips.2009.04.001

18. Scarselli M, Donaldson JG. Constitutive Internalization of G Protein-coupled Receptors and G Proteins via Clathrin-independent Endocytosis. Journal of Biological Chemistry. 2009;284: 3577–3585. doi:10.1074/jbc.M806819200

19. Kirkham M, Parton RG. Clathrin-independent endocytosis: New insights into caveolae and non-caveolar lipid raft carriers. Biochimica et Biophysica Acta (BBA) - Molecular Cell Research. 2005;1745: 273–286. doi:10.1016/j.bbamcr.2005.06.002

20. Martinez-Outschoorn UE, Sotgia F, Lisanti MP. Caveolae and signalling in cancer. Nat Rev Cancer. 2015;15: 225–237. doi:10.1038/nrc3915

21. Chisari M, Saini DK, Kalyanaraman V, Gautam N. Shuttling of G Protein Subunits between the Plasma Membrane and Intracellular Membranes. Journal of Biological Chemistry. 2007;282: 24092–24098. doi:10.1074/jbc.M704246200

22. Calebiro D, Koszegi Z, Lanoiselée Y, Miljus T, O’Brien S. G protein-coupled receptor-G protein interactions: a single-molecule perspective. Physiol Rev. 2021;101: 857–906. doi:10.1152/physrev.00021.2020

23. Bondar A, Jang W, Sviridova E, Lambert NA. Components of the Gs signaling cascade exhibit distinct changes in mobility and membrane domain localization upon β2-adrenergic receptor activation. Traffic. 2020;21: 324–332. doi:10.1111/tra.12724

24. van Hemert F, Lazova MD, Snaar-Jagaska BE, Schmidt T. Mobility of G proteins is heterogeneous and polarized during chemotaxis. J Cell Sci. 2010;123: 2922–2930. doi:10.1242/jcs.063990

25. Sungkaworn T, Jobin M-L, Burnecki K, Weron A, Lohse MJ, Calebiro D. Single-molecule imaging reveals receptor–G protein interactions at cell surface hot spots. Nature. 2017;550: 543–547. doi:10.1038/nature24264

26. Qin K, Dong C, Wu G, Lambert NA. Inactive-state preassembly of G(q)-coupled receptors and G(q) heterotrimers. Nat Chem Biol. 2011;7: 740–747. doi:10.1038/nchembio.642

27. Jang W, Adams CE, Liu H, Zhang C, Levy FO, Andressen KW, et al. An inactive receptor-G protein complex maintains the dynamic range of agonist-induced signaling. Proc Natl Acad Sci U S A. 2020;117: 30755–30762. doi:10.1073/pnas.2010801117

28. Petelák A, Lambert NA, Bondar A. Serotonin 5-HT7 receptor slows down the Gs protein: a single molecule perspective. MBoC. 2023;34. doi:10.1091/mbc.E23-03-0117

29. Yanagawa M, Hiroshima M, Togashi Y, Abe M, Yamashita T, Shichida Y, et al. Single-molecule diffusion-based estimation of ligand effects on G protein–coupled receptors. Science Signaling. 2018;11: eaao1917. doi:10.1126/scisignal.aao1917

30. Mystek P, Tworzydło M, Dziedzicka-Wasylewska M, Polit A. New insights into the model of dopamine D1 receptor and G-proteins interactions. Biochimica et Biophysica Acta (BBA) - Molecular Cell Research. 2015;1853: 594–603. doi:10.1016/j.bbamcr.2014.12.015

31. Gibson SK, Gilman AG. Giα and Gβ subunits both define selectivity of G protein activation by α2-adrenergic receptors. Proceedings of the National Academy of Sciences. 2006;103: 212–217. doi:10.1073/pnas.0509763102

32. Bondar A, Lazar J. Dissociated GαGTP and Gβγ Protein Subunits Are the Major Activated Form of Heterotrimeric Gi/o Proteins. Journal of Biological Chemistry. 2014;289: 1271–1281. doi:10.1074/jbc.M113.493643

33. Ershov D, Phan M-S, Pylvänäinen JW, Rigaud SU, Le Blanc L, Charles-Orszag A, et al. TrackMate 7: integrating state-of-the-art segmentation algorithms into tracking pipelines. Nat Methods. 2022;19: 829–832. doi:10.1038/s41592-022-01507-1

34. Vega AR, Freeman SA, Grinstein S, Jaqaman K. Multistep Track Segmentation and Motion Classification for Transient Mobility Analysis. Biophys J. 2018;114: 1018–1025. doi:10.1016/j.bpj.2018.01.012

35. Nobles M, Benians A, Tinker A. Heterotrimeric G proteins precouple with G protein-coupled receptors in living cells. Proceedings of the National Academy of Sciences. 2005;102: 18706–18711. doi:10.1073/pnas.0504778102

36. Kleuss C, Krause E. Gαs is palmitoylated at the N-terminal glycine. EMBO J. 2003;22: 826–832. doi:10.1093/emboj/cdg095

37. Chytła A, Gajdzik-Nowak W, Biernatowska A, Sikorski AF, Czogalla A. High-Level Expression of Palmitoylated MPP1 Recombinant Protein in Mammalian Cells. Membranes. 2021;11: 715. doi:10.3390/membranes11090715

38. Kosloff M, Elia N, Selinger Z. Structural Homology Discloses a Bifunctional Structural Motif at the N-Termini of Gα Proteins. Biochemistry. 2002;41: 14518–14523. doi:10.1021/bi026729x

39. Mystek P, Martikainen A, Błasiak E, Kulig W, Polit A. The polybasic puzzle: how N-terminal charges modulate Gαi1 membrane behavior and signaling output. Cell Commun Signal. 2025;23: 547. doi:10.1186/s12964-025-02541-0

