## Supporting information for "Heterotrimeric G proteins containing Gβ_1_γ_2_ subunits exhibit subtype-specific mobility differences in live cells"

### Supporting information for “Heterotrimeric G proteins containing $G\beta_1\gamma_2$ subunits exhibit subtype-specific mobility differences in live cells”

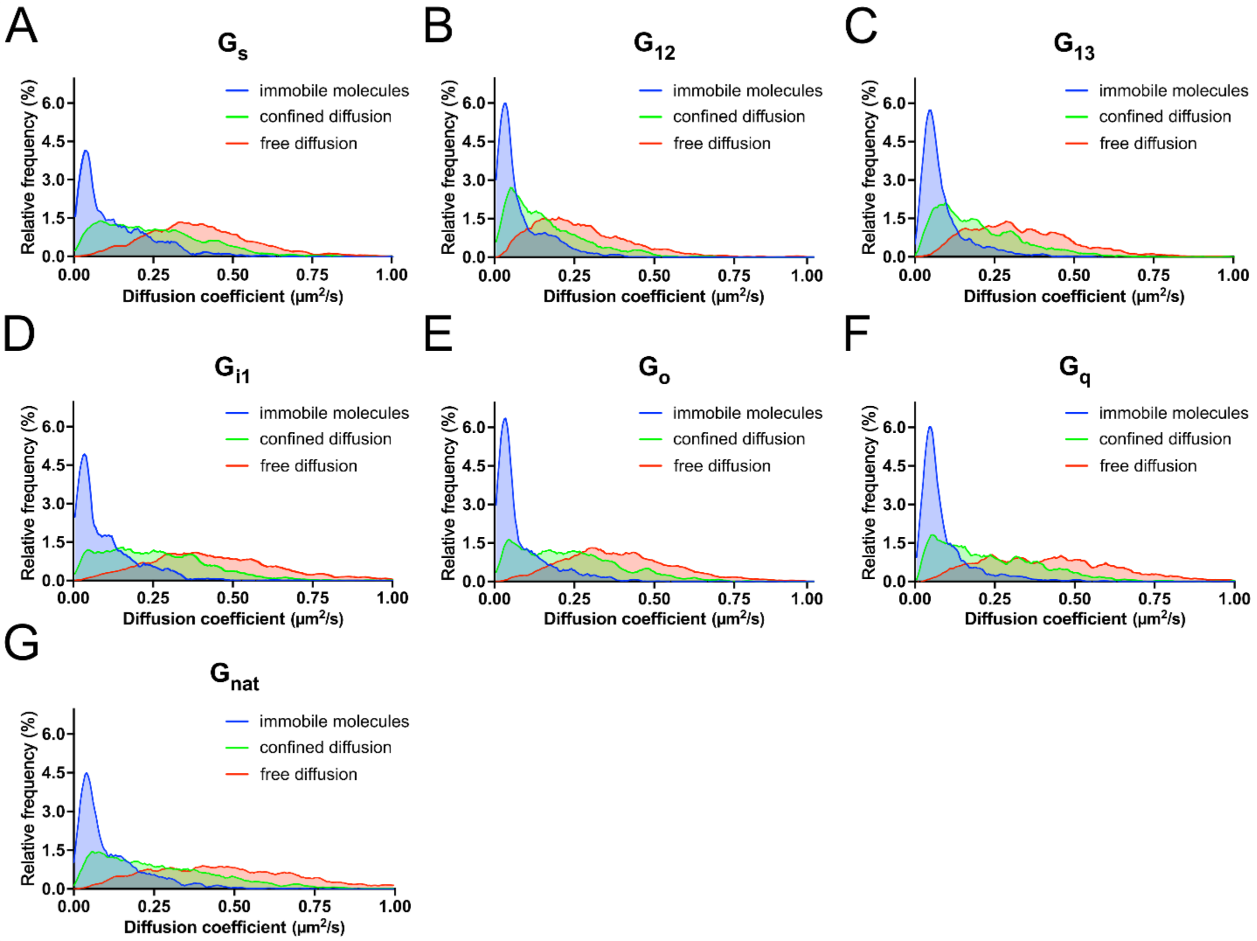


**S1 Fig. Distribution of diffusion states in G proteins.** Graphical comparison of the distribution of diffusion coefficients of molecular tracks based on diffusion state — immobile (blue), confined (green), freely diffusing (red), in $G_s$ (A), $G_{12}$ (B), $G_{13}$ (C), $G_{i1}$ (D), $G_o$ (E), $G_q$ (F), and $G_{nat}$ (G). Molecule tracks exhibiting directed diffusion are not shown in this representation due to very low sample size. No statistical inference was performed on these distributions.

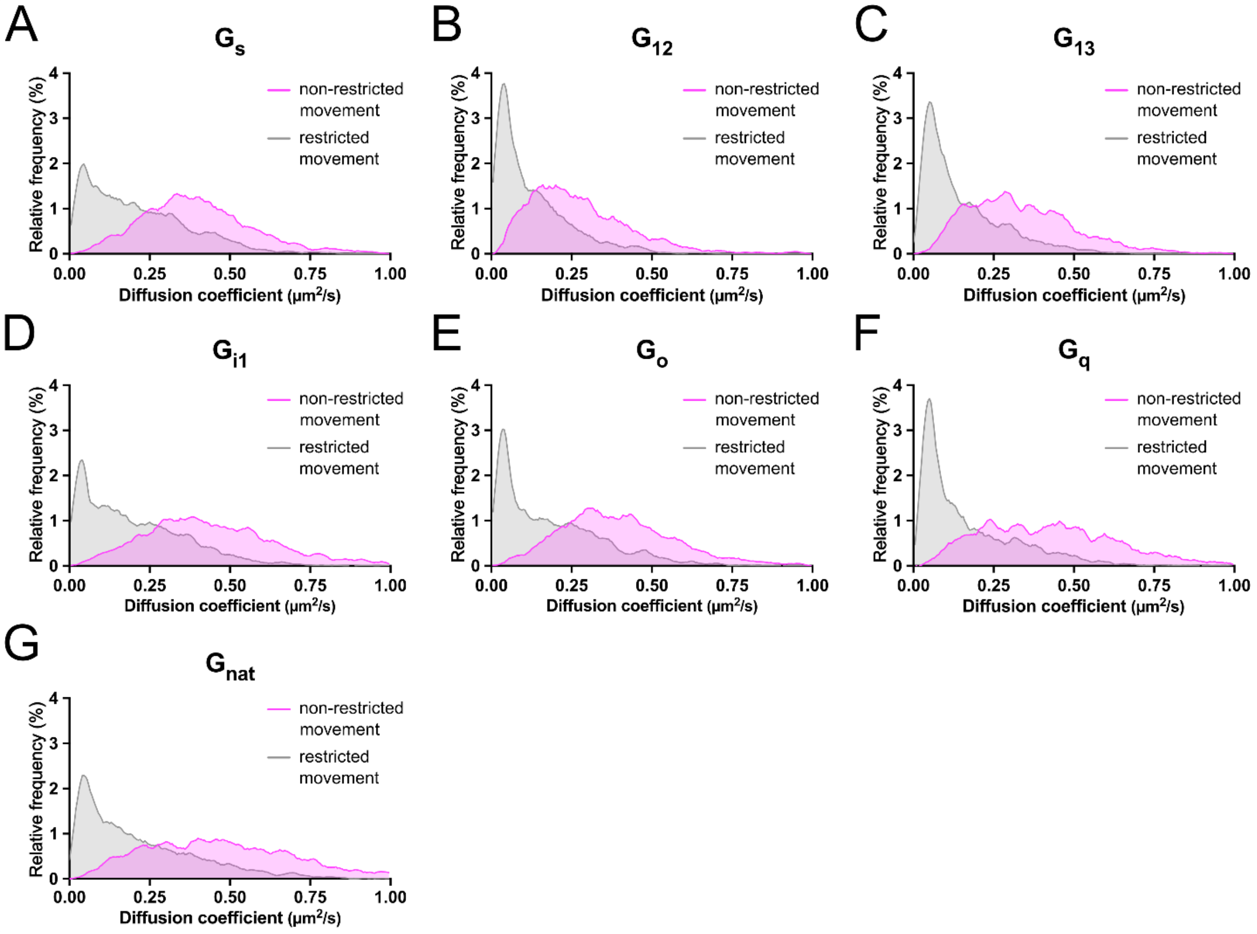


**S2 Fig. Distribution of restricted and non-restricted diffusion types in G proteins.** Graphical comparison of the distribution of diffusion coefficients of restricted (grey) and non-restricted molecule tracks (magenta) of $G_s$ (A), $G_{12}$ (B), $G_{13}$ (C), $G_{i1}$ (D), $G_o$ (E), $G_q$ (F), and $G_{nat}$ (G). No statistical inference was performed on these distributions.

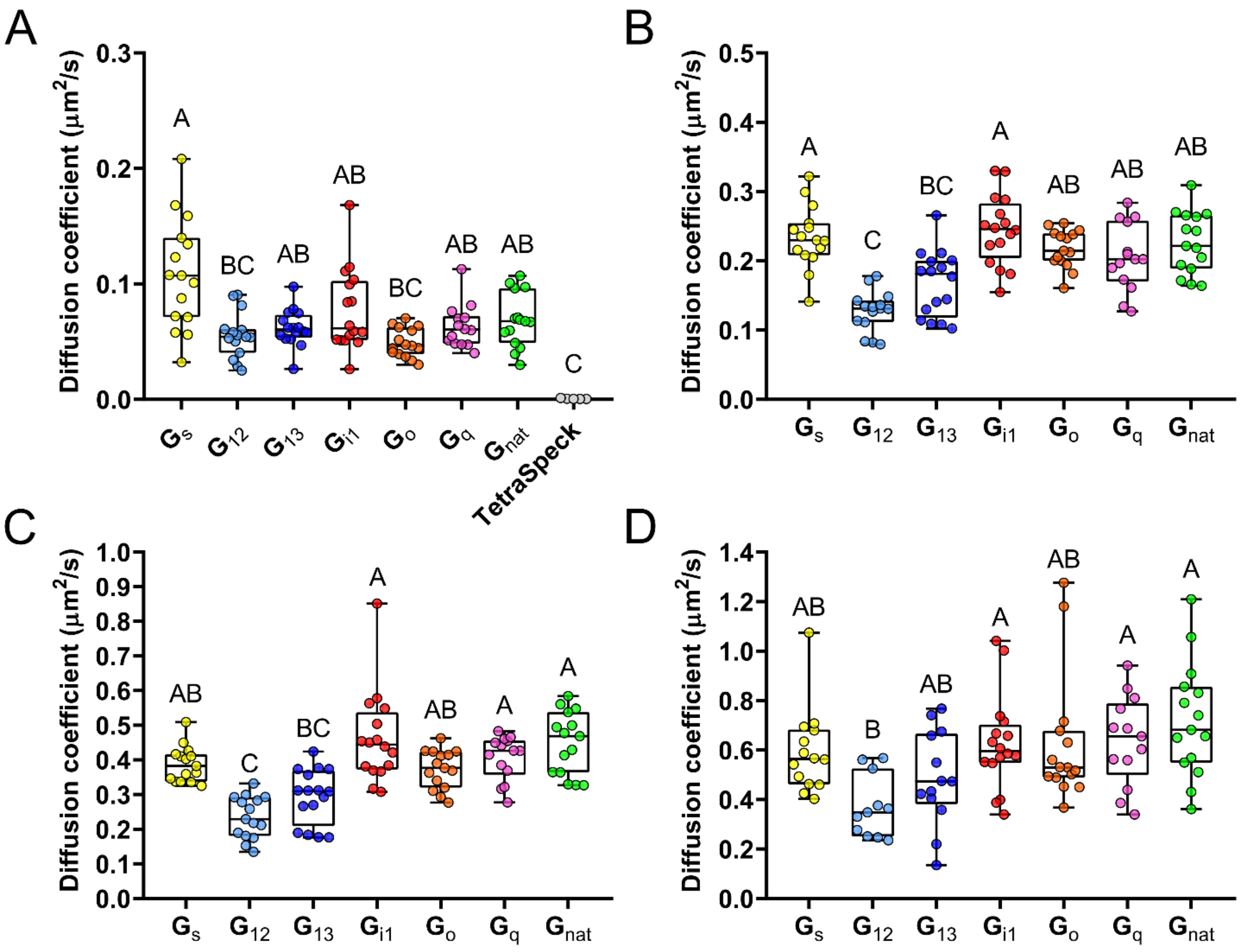


**S3 Fig. G proteins show diffusion state-specific differences in mobility.** A-D) Box plots of median diffusion coefficients of immobile molecules (A) and molecule tracks exhibiting confined (B), free (C), and directed diffusion (D). The boxes indicate the median and the interquartile range. Compact letter display above individual datasets indicates similarity grouping based on statistical analysis ($\alpha = 0.05$; A: $p < 0.0001$, $H = 36.65$, $\varepsilon^2 = 0.29$; B: $p < 0.0001$, $H = 46.25$, $\varepsilon^2 = 0.41$; C: $p < 0.0001$, $H = 54.43$, $\varepsilon^2 = 0.49$; D: $p < 0.0001$, $H = 23.65$, $\varepsilon^2 = 0.20$).

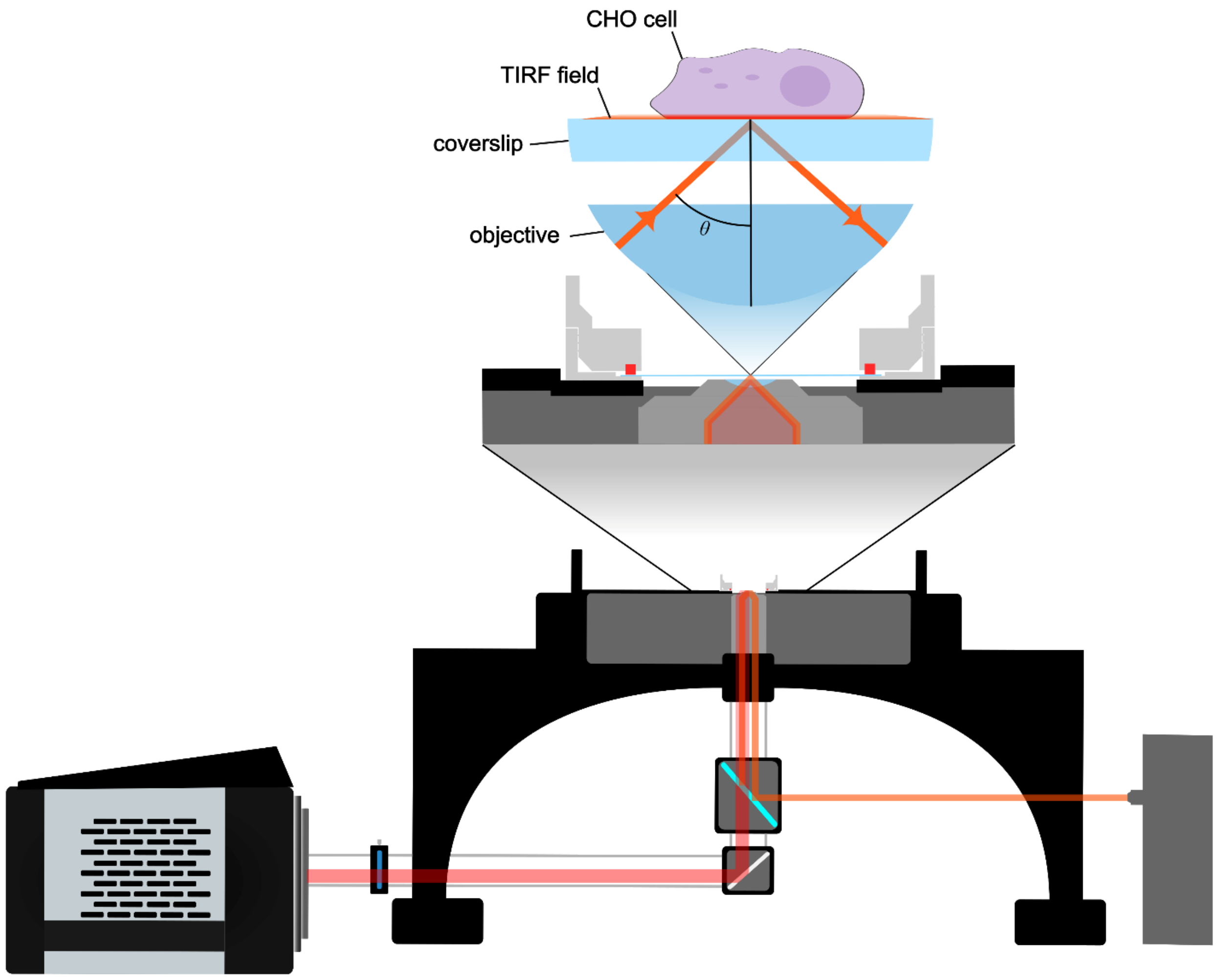


**S4 Fig. Single-molecule TIRF microscopy setup.** An inverted custom-built TIRF microscope used for single-molecule imaging utilizing a 60× 1.49NA objective lens, a 640 nm laser, and an sCMOS Sona camera. The top diagram shows a close-up of the sample illumination achieved during TIRF microscopy.

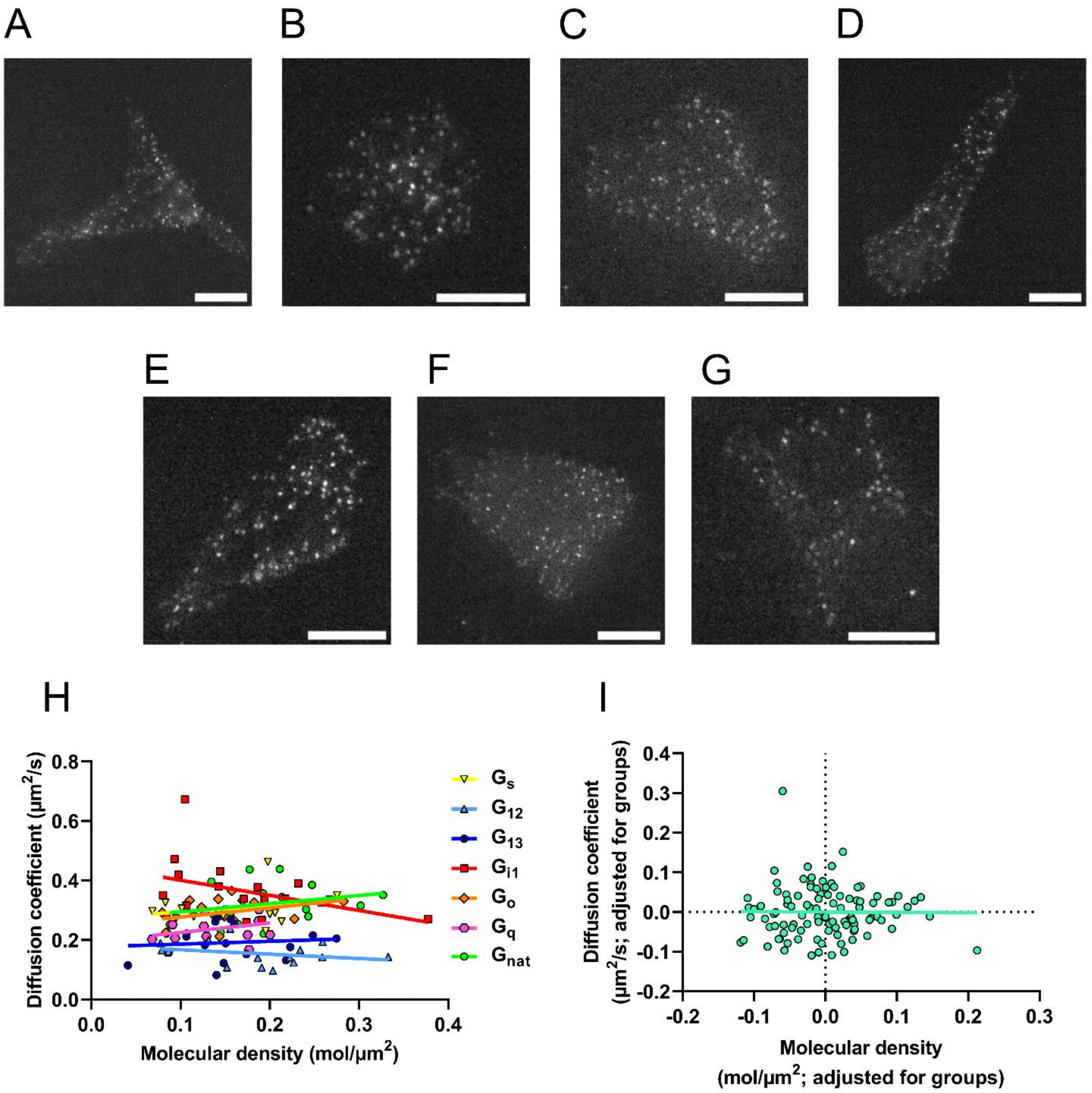


**S5 Fig. Membrane density and mobility in G protein heterotrimers.** Representative images of $G_s$ (A), $G_{12}$ (B), $G_{13}$ (C), $G_{i1}$ (D), $G_o$ (E), $G_q$ (F), and $G_{nat}$ (G) expressing cells. The scale bar in A-G is 10 μm. H) Linear regression fit illustrating the relationship between the mobility and membrane density of labeled $G\gamma_2$-containing heterotrimers. Due to low sample size, no statistical inference was performed on these values. I) Linear regression analysis of the correlation between the mobility and membrane density of labeled $G\gamma_2$-containing heterotrimers adjusted for group identity through residual analysis (slope = -0.009, 95% CI: −0.203 to 0.186; $R^2$ = <0.001; $F(1, 104) = 0.008$; $p = 0.928$; $n = 106$).

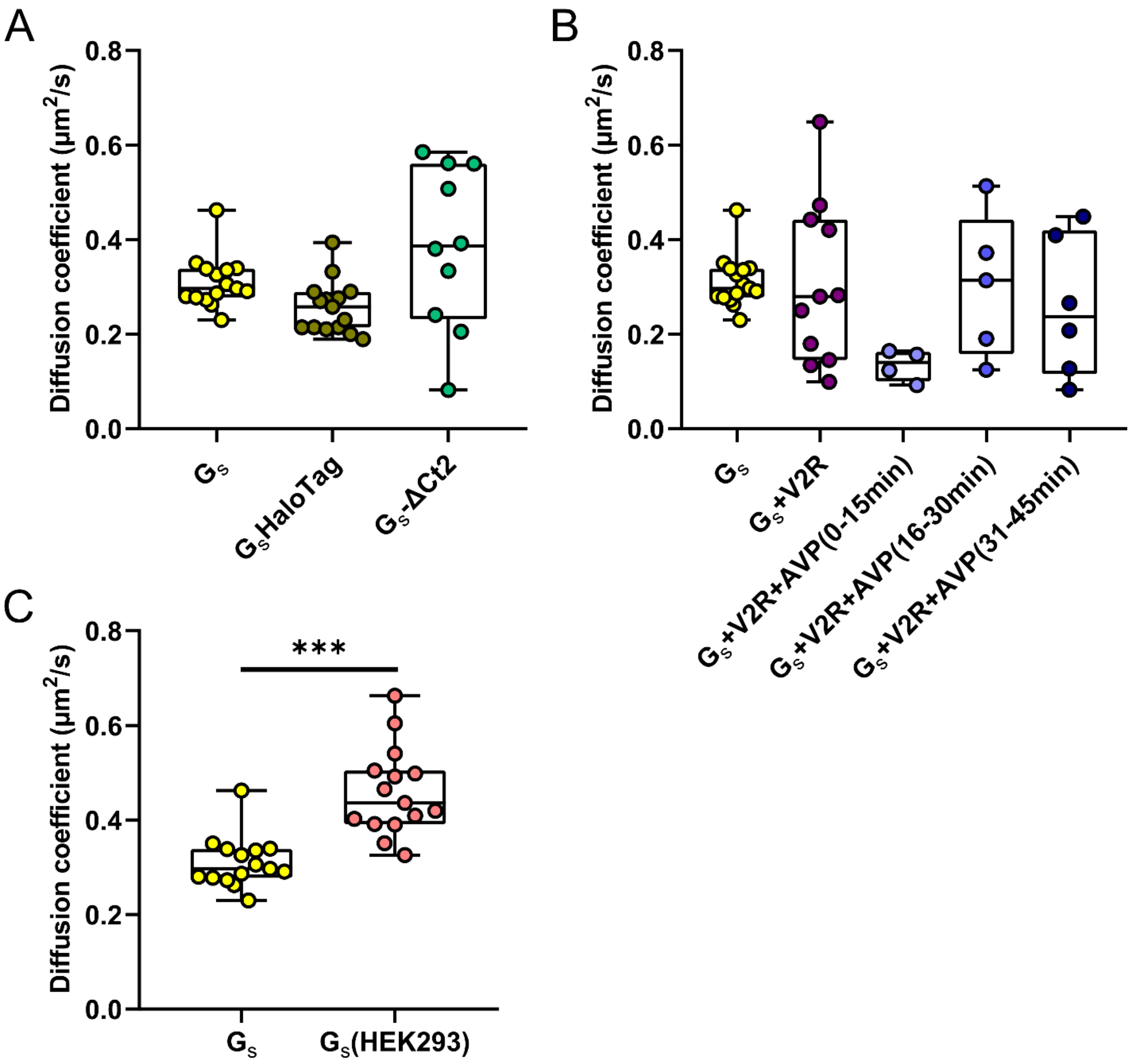


**S6 Fig. Effect of labeling and experimental system on the $G_s$ protein mobility.** A) Median diffusion coefficients of $G_s$ heterotrimers observed using overexpression of $G\alpha_s$ and $G\beta_1$ under the CMV promoter, and underexpression of $G\gamma_2$-HaloTag under the minP promoter ($G_s$: *n* = 15, observed range = 0.23-0.46 μm²/sec), as well as through overexpression of $G\gamma_2$ and $G\beta_1$ under the CMV promoter, and underexpression of $G\alpha_s$-HaloTag under the minP promoter ($G_s$HaloTag; *n* = 15, observed range = 0.19-39 μm²/sec). Additionally, diffusion coefficient medians of $G\gamma_2$-labeled heterotrimers containing the $G\alpha_s$-ΔCt2 mutant, missing two C-terminal amino acids, are shown ($G_s$-ΔCt2; *n* = 10, observed range = 0.08-0.58 μm²/sec). B) Diffusion coefficient medians of the $G_s$ protein (*n* = 15, observed range = 0.23-0.46 μm²/sec) coexpressed with the vasopressin 2 receptor ($G_s$+V2R, untreated; $n$ = 11, observed range = 0.1-0.64 μm$^2$/sec) and treated with arginine vasopressin 1μM ($G_s$+V2R+AVP, separated into time intervals post-treatment; 0-15 min: $n$ = 4, observed range = 0.09-0.16 μm$^2$/sec; 16-30 min: $n$ = 5, observed range = 0.13-0.51 μm$^2$/sec; 31-45 min: $n$ = 6, observed range = 0.08-0.44 μm$^2$/sec). The boxes indicate the median and the interquartile range. Due to low sample size, no statistical inference was performed on the medians shown in A and B. C) Diffusion coefficient medians of $G_s$ expressed in CHO-K1 cells ($G_s$) and HEK-293 cells ($G_s$(HEK-293)). Statistical analysis results are indicated above ($\alpha$ = 0.05, $p$ < 0.0001, $U$ = 12, $r_{rb}$ = 0.89).

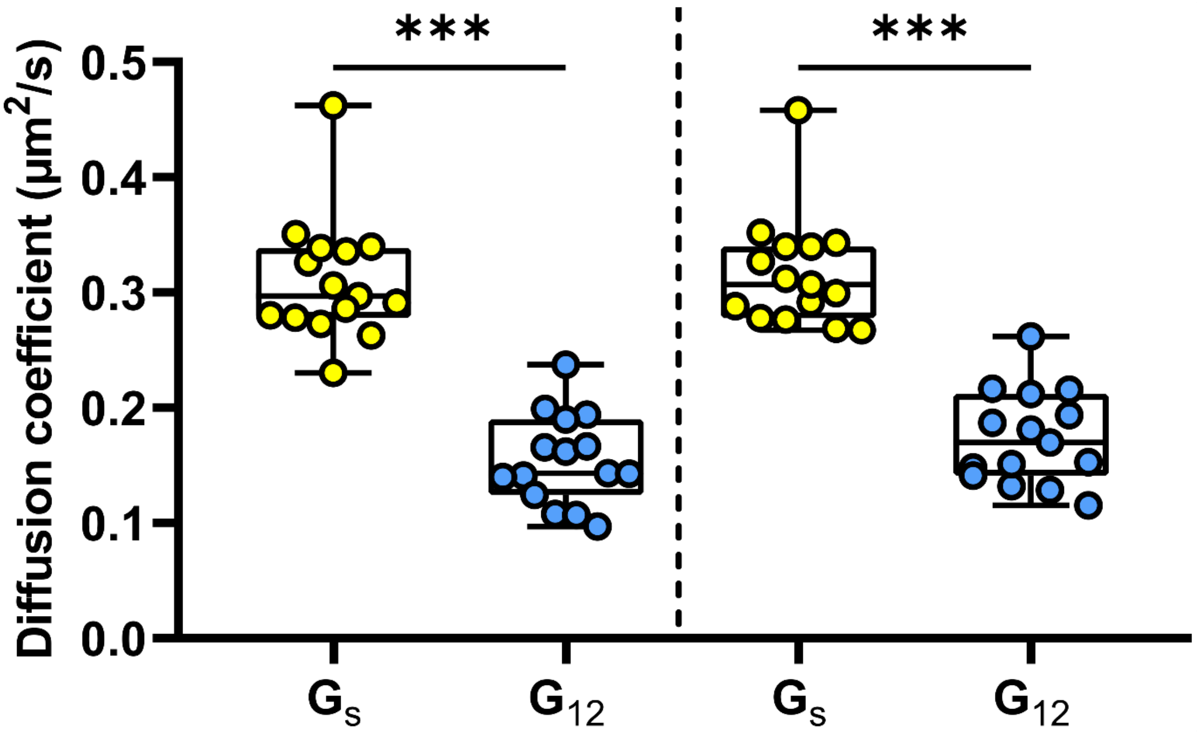


**S7 Fig. Comparison of alternative data analysis approach results.** On the left side, diffusion coefficient medians of $G_s$ and $G_{12}$ obtained through analysis using input parameters adjusted on a case-by-case basis as previously indicated. Statistical analysis results are indicated above ($\alpha$ = 0.05, $p$ < 0.0001, $U$ = 1, $r_{rb}$ = 0.99). On the right side, diffusion coefficient medians of $G_s$ and $G_{12}$ obtained through an alternative analysis approach using fixed input parameters ($\alpha$ = 0.05, $p$ < 0.0001, $U$ = 0, $r_{rb}$ = 1).

| | $G_s$ | $G_{12}$ | $G_{13}$ | $G_{i1}$ | $G_o$ | $G_q$ | $G_{nat}$ |
|---|---|---|---|---|---|---|---|
| Molecular density (molecule/μm$^2$) ± SD | 0.173 ±0.064 | 0.187 ± 0.071 | 0.160 ± 0.056 | 0.165 ± 0.076 | 0.149 ± 0.052 | 0.139 ± 0.041 | 0.202 ± 0.061 |

**S1 Table. Membrane density of the labeled G proteins.** The membrane density of G$\gamma_2$-labeled heterotrimers based on the number of molecules detected in the first frame of each image series.

| | Number of separate transfection experiments | Number of analyzed cells | Number of analyzed tracks |
|---|---|---|---|
| $G_s$ | 3 | 15 | 7129 |
| $G_{12}$ | 4 | 15 | 5278 |
| $G_{13}$ | 4 | 16 | 4805 |
| $G_{i1}$ | 3 | 16 | 6190 |
| $G_o$ | 3 | 15 | 5341 |
| $G_q$ | 3 | 14 | 3944 |
| $G_{nat}$ | 3 | 15 | 7543 |

**S2 Table. Data summary.** The table shows the number of separate transfection experiments, cells, and tracks analyzed for each condition.
